# Gut cancer increases the risk for *Drosophila* to be preyed upon by hunting spiders

**DOI:** 10.1101/2020.07.01.182824

**Authors:** David Duneau, Nicolas Buchon

## Abstract

Predators are thought to prey on individuals that are in poor physical condition, although the evidence supporting this is ambiguous. We tested if sick individuals where more predated using *Drosophila melanogaster* flies as manipulable preys. We asked whether hunting spiders, trapped from the wild, would selectively prey upon flies with compromised health (i.e. chronically infected or cancerous) versus healthy flies under laboratory conditions. Flies chronically infected with the bacterium *Providencia rettgeri*, a natural *Drosophila* pathogen, were not selectively preyed upon by jumping spiders. We strengthened and confirmed our finding with another hunting spider species, small wolf spiders. We discuss that this result supports the hypothesis that chronic infection is associated with reduced symptoms notably to avoid the potentially deadly consequences of host predation on pathogens. We then induced colon cancer in some of the flies and asked whether the presence of cancer led to selective predation; there is little evidence for this, even in vertebrates. As the cancer developed, the incidence of predation by jumping spiders on the afflicted flies increased. We conclude that disease can have different lethal consequences through predation, even in invertebrate species, and that cancer is a factor in selective predation. Our results may explain why early tumours, but not metastasized cancers, are commonly detected in organisms in the wild, as cancer bearing individuals are rapidly eliminated due to the strong selective pressure against them.

## Introduction

In vertebrates, individuals that are injured, or diseased are often preyed upon, as are juveniles who have not reached adult speed or reflex (Furey et al., 2021; Genovart et al., 2010; Mesa, Poe, Gadomski, & Petersen, 1994; Møller, 2008; Murray, 2002). This is especially true when a predator’s preferred type of prey is difficult to catch at the prime of its performance (Temple, 1987), however, there are exceptions (Penteriani et al., 2008). Although this conventional wisdom of selective predation on substandard individuals is largely accepted, only a few studies have tested it empirically. Selective predation can only be said to exist when the relative frequencies of the types of prey in a predator’s diet differ from the frequencies of the prey in the environment (Chesson, 1978). Hence, to test the selective prey hypothesis in the wild, data on the types of prey sought by many of the predators and the health status of the whole prey population would be determined. This is obviously difficult to obtain. Furthermore, a prey individual may suffer from different conditions which influence each other and lead to spurious effects. For example, exhaustion from care for large brood increases susceptibility to parasites (Oppliger, Christe, & Richner, 1996). Thus, the fittest hosts are the ones classified as being sick. Although those information may be vitally important in the context of eco-evolutionary dynamics (Brunner, Anaya-Rojas, Matthews, & Eizaguirre, 2017; Brunner, Deere, Egas, Eizaguirre, & Raeymaekers, 2019), the topic has been subjected more to speculation than to empirical study.

Probably the most studied question regarding selective predation is whether predators prey more on individuals with suboptimal conditions over healthy individuals. Even if hosts predators may avoid infected prey to avoid the risk of getting sick themselves (Goren & Ben-Ami, 2017), it has been suggested, based on field observations, that infections could increase the vulnerability to predation (Adelman, Mayer, & Hawley, 2017; Duffy, Hall, Tessier, & Huebner, 2005; Gooding et al., 2020; Miller et al., 2008; Moller & Erritzoe, 2000; Moller, Erritzoe, & Tottrup, 2010; Møller & Nielsen, 2007; Murray, Cary, & Keith, 2006) and there is some experimental support for this (DeBlieux & Hoverman, 2019; Gallagher et al., 2019; Johnson, Stanton, Preu, Forshay, & Carpenter, 2006; Murray, 2002). Consequently, by mediating selective predation, parasites can mediate the relationship between hosts and their prey (Hall, Cáceres, Duffy, & Cáceres, 2005; Møller, 2008). For instance, removing predators can reduce vertebrate prey populations (Sih, Crowley, Mcpeek, Petranka, & Strohmeier, 1985), which seems counterintuitive, as predators are expected to remove preys. However, mathematical models predict that virulent parasites may be selected against if predation disproportionately removes them before they can be transmitted to the host (Packer, Holt, Hudson, Lafferty, & Dobson, 2003). Without this check on virulent parasites by the predator, more prey may die from lethal infections than from predation. Hence, if selective predation for infected individuals is a common pattern, it may also be a major driver of host and parasite evolution and cannot be ignored any longer (Møller, 2008).

It is also relevant to consider that not all sickness come from infections and that non-infectious diseases could also increase the risk of being predated. In fact, understanding the role of cancer on ecosystem function has been identified as a key endeavour (Dujon, Aktipis, et al., 2021). So far, most of our knowledge is based on the consequence of transmissible cancer on an apex predator (i.e. Tasmanian devil (Cunningham, Johnson, & Jones, 2020; Hollings, Jones, Mooney, & Mccallum, 2014; Hollings, Jones, Mooney, & McCallum, 2016; Woods et al., 2018)), and few to nothing is known about the potential selective predation of prey that are sick due to a non-infectious disease (see (Perret, Gidoin, Ujvari, Thomas, & Roche, 2020) for a theoretical study). Mutations can occur during the cell replication and division required to grow and maintain multicellular animals, and certain types of mutation, said oncogenic mutations, can lead to the formation of tumours (Aktipis et al., 2015; Albuquerque, Drummond do Val, Doherty, & de Magalhães, 2018; Hanahan & Weinberg, 2011; National Cancer Institute, 2020a). In most cases, tumours are benign (Bissell & Hines, 2011), but they can be malignant, and even fatal (Aktipis et al., 2015; Albuquerque et al., 2018; Hanahan & Weinberg, 2011; National Cancer Institute, 2020b). Clinical categorization of tumours has been established in unchallenging, even ‘optimal’ environments (i.e. in the laboratory or clinic), usually to the point of death via organ failure. The impact of tumour progression is likely to influence tremendously individual’s life history traits (e.g. lifespan and reproductive output but also competitive and dispersal ability, and pathogen susceptibility), yet, its impact under more natural conditions has largely been neglected (Roche, Møller, DeGregori, & Thomas, 2017; Marion Vittecoq et al., 2013). Furthermore, depending on the relationship between tumour development and the health of the prey, predation could affect the selection of oncogenic mutations and thus the risk to develop cancer. To our knowledge, the possible interactions between a predator and a prey that is afflicted with cancer are poorly understood and have not been empirically studied to date (Marion Vittecoq et al., 2013).

Here, we induced chronic bacterial infection and colon cancer in prey to understand their role in increasing the likelihood of being predated. We use the fruit fly *Drosophila melanogaster,* a model which has started recently to be used to study the effect of cancer from an ecological perspective (Arnal et al., 2017; Dawson et al., 2018) and is already a well-characterized model for infectious and non-infectious diseases, as a novel model for sick prey. We assess the predation of flies by hunting spiders, using flies chronically infected by a wild-caught bacterial pathogen *(Providencia rettgeri),* and have genetically induced cancer, under laboratory-controlled conditions. These conditions allow for the ideal experimental setting to study predation and the role of prey well-being in a population context.

## Materials and methods

### Prey and predators

We used *Drosophila melanogaster* flies as prey in this study as they are tractable to genetic induction of tumours, are well-defined models for bacterial infection, and allow for control of environmental and genetic differences. Only one sex (males) were used, to avoid sex effects. We induced chronic infection by injecting 23 nL of a suspension of few thousand cells of the bacterium *Providencia rettgeri* (strain *Dmel*, isolated from wild-caught *Drosophila melanogaster*; Juneja and Lazzaro 2009) in phosphate-buffered saline (PBS) into the abdomen of flies (Canton S line, a wildtype genotype kept in the laboratory since several decades). The same volume of sterile PBS without bacteria was injected into another group of flies that served as the control. Individuals surviving for three days or more after injection of the bacteria have an established chronic infection, as we have previously demonstrated (Duneau et al., 2017).

To induce tumours in the flies, we took advantage of signalling pathways that regulate cell growth in mammals, which have a conserved function in *Drosophila*, the EGFR and Wnt pathways (Millburn et al., 2016; Mirzoyan et al., 2019; Rudrapatna, Cagan, & Das, 2012; Villegas, 2019). We used targeted induction of these pathways (Rasv12 for EGFR, APC RNAi for the Wnt pathway) to induce two types of enlargement of the flies digestive tract (i.e. hyperplasia, often called benign tumour, and cancer). Those types of colon cancer is appropriate for this study, as it does not overtly affect physical performance including locomotion, and will thus allow for the study of the interaction of oncogenic phenomena and predation (see Dawson et al. 2018 for locomotion assays in gut cancerous flies in another genetic background). To generate tumours in the gut of *Drosophila*, flies *esg^TS^* (*esgGa14, Ga18^ts^, UAS-GFP*) were crossed to either background control flies (Trip stock 35786), flies carrying a *UAS-Ras^v12^* (Bloomington 64196) or flies carrying a combination of *UAS-Ras^v12^* (Bloomington 641926) and *UAS-APC-RNAi* (Bloomington Trip stock 28582) combined. The expression of Ras^v12^ in progenitor cells stimulates an accelerated turnover and a mild accumulation of epithelial cells in the gut, akin to a dysplastic tissue (Buchon, Broderick, Kuraishi, & Lemaitre, 2010; Houtz, Bonfini, Bing, & Buchon, 2019), later also referred as hyperplasia. The overexpression of *Ras^v12^* combined with the knock-down of *APC* leads to increased proliferation, and accumulation of progenitor cells and epithelial cells in the gut, reminiscent of an intestinal disseminated tumour (Wang et al., 2013), later also referred as cancerous. F1 flies from these crosses were raised at 18°C (Gal4 off, transgene expression off, normal development and emergence), and 3 days after emergence were switched to 29°C (Gal4 on, transgene expression on) for 10 or 20 days before performing the experiment. As in (Wang et al., 2013), crosses with WT flies showed mild tissue renewal (restricted GFP signal), flies with expressing *Ras^v12^* showed disseminated GFP in their gut and flies expressing *Ras^v12^* and *APC RNAi* showed gut enlargement and GFP accumulation. The longest the induction is done and the strong is the phenotypes leading to a gradient from induced hyperplasia for 10 days (i.e. mild colon enlargement) to induced cancer for 20 days (i.e. limit to the complete obstruction of the colon).

We used two families of hunting spiders as predators: jumping spiders of several species from the family of Salticidae (see illustration in Fig. 1) and small wolf spiders from the family of Lycosidae (likely *Pardosa lugubris*). Both jumping and wolf spiders hunt by wandering rather than by using webs as traps.

**Figure 1.**
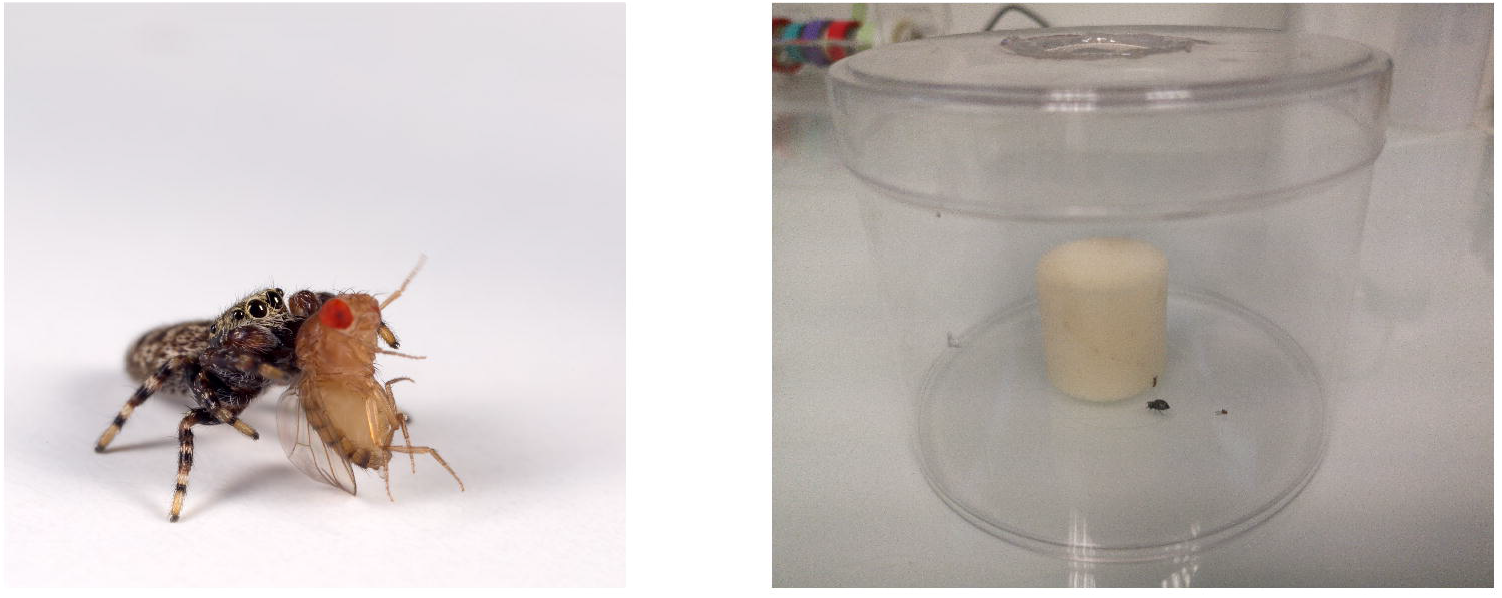
Photograph of a jumping spider consuming *Drosophila melanogaster* flies in the laboratory.

Jumping spiders have the sharpest vision of any arthropod, even surpassing many vertebrates (Land & Nilsson, 2012). They hunt during the day and rely on this astonishing vision when catching a prey. Similar to the way a cat stalks its prey, jumping spiders turn towards their prey, directed by a pair of lateral eyes that provide a nearly panoramic field of view with the ability to discern motion (Zurek & Nelson, 2012). Then, the spiders track, approach, and jump on the prey. The vision accuracy is conferred by two pairs of forward-facing eyes that also provide a precise perception of depth (Nagata et al., 2012; Zurek, Taylor, Evans, & Nelson, 2010).

After having tested jumping spiders as predator, we decided to test whether infected individuals would still not be more likely to be eaten by another type of hunting spider species with different hunting behaviour, wolf spiders. This was additionally performed to strengthen our “negative” result. Wolf and jumping spiders have already successfully been used to test for selective predation (Holmberg & Turnbull, 1982; Vickers & Taylor, 2018). Wolf spiders are generally nocturnal hunters and their eyes function mainly as low-light movement detectors, however *Pardosa* species are mostly hunting during the day (Edgar, 1969). Their vision is not as acute as that of jumping spiders (Land & Nilsson, 2012). The use of visual cues in prey detection and orientation have only been well-studied in jumping spiders, and empirical evidence suggests that wolf spiders rely more on vibrations to capture their prey than on vision (Lizotte & Rovner, 1988). Hence, the differences between these two types of hunting spiders primarily lie in their ability to prey on flies that are at rest. In an environment where there is no place to hide, jumping spiders would likely prey equally well on moving and non-moving prey, and selective predation in this case should depend largely on the prey’s capacity to escape.

### Predation trials

Predation trials were performed during the day by incubating five healthy and five sick flies, age-matched, at 20°C with a spider in a round plastic box (11.5cm × 8.5cm, see illustration in fig. 1B) containing a wet piece of cotton or foam. The flies were first added to the box to settle before the introduction of the predators. After about 30 min, spiders were put in the box. The trials generally lasted for about 4 hours or were stopped when 50% of the flies had been preyed upon, as, when scoring only at the end of the experiment, it allows to score the most extreme result (i.e. all individuals from one treatment have been eaten before the other treatment). Even if no spiders were starved before the experiment, this goal was sometimes exceeded as some spiders ate faster than expected. Individual spiders have never been used twice as predators within an experiment.

To determine the number of surviving flies that were infected, we ground the remaining flies at the end of the predation period and suspended the mixture in 250 μL of Luria-Bertani (LB) medium. A droplet of five microliters of the resulting suspension was spread on a petri dish filled with agar containing LB medium, and the plates were incubated overnight at 37°C to assess for the presence of *P. rettgeri*. We calculated the proportion of infected flies at the end of predation period by determining the number of droplets in which *P. rettgeri* were present. For a control comparison, we performed the same process with infected and healthy flies, but did not expose them to predatory spiders. Our protocol consistently led to the induction of tumour, even if interindividual variation in tumour size exist which can be a source of variation in our trials. In the unlikely event that some individuals were considered has carrying a tumour but did not, those individuals would reduce the strength of our signal as they would be like control individuals. The number of surviving flies that had cancer were counted by dissecting all surviving flies and observing their guts under a fluorescent microscope, which reveals the presence/absence of cancer.

### Statistical analysis

Statistical analysis were performed using R, version 4.1 (R Core Team, 2020). The use of the Fisher exact test or a chi-squared test would have permitted us to test for every trial regardless of whether the number of flies from a particular category was more or less preyed upon than those from another category. However, we used a hypergeometric distribution instead of a Chi-squared distribution because, as flies were preyed upon, the population size decreased and we only had information on predation rate per treatment at the end of the experiment. The *p-*value for each experimental trial was added up and compared to the summed *p*-value obtained by simulating random trials. We assessed the overall *p*-value by determining the number of simulated trials required to have an observed result (summed *p*-value observed) as probable as the simulation (summed *p*-value simulated). Hence, a *p*-value of 0.05 indicated that 20 trials would be sufficient to have the same result as the empirical data by chance, whereas a *p*-value of 0.0001 meant that 10,000 trials would be required to achieve the same result as the empirical data by chance. We used the Manly Alpha Index as the preference index to illustrate the selective predation on sick individuals (index 0: preference for healthy individuals, index 0.5: no preference, index 1: preference for sick individuals). It was calculated as log (total flies sick / flies sick but not eaten) / [(log(total flies sick / flies sick but not eaten) + log(total flies healthy / flies healthy but not eaten)] as established by Manly (1972). To test whether the preference index for sick individuals increased with the severity of the cancer, we considered hyperplasia after 10 and 20 days of induction and cancer after 10 days and 20 days as ordinal variables. This was used because although we knew the rank of severity (i.e. from inducing hyperplasia for 10 days to inducing cancer for 20 days), we could not know the size of the differences among them. This allowed us to test in a single model (*i.e*. using an ordinal logistic regression) for the increase of the preference index with the increase in cancer severity.

### Ethical note

All spiders in the study were obtained from the wild during their daily activity and kept for less than a week, eventually fed with healthy *Drosophila*. Spiders used for testing the effect of cancer were sampled and tested on the Cornell University campus (Ithaca, NY, USA) and those used for testing the effect of infectious diseases were sampled and tested on the campus of the University of Toulouse 3 (Toulouse, France). All individuals have been released in the wild at the area they have been caught shortly after the experiments were performed.

## Results

### Predation of chronically infected flies by hunting spiders

We tested whether chronic infection of the male flies by *P. rettgeri* affected the risk of being preyed upon by the spiders. The flies had been infected for several days before exposure to the spiders, and their immune response was expected to be activated (Chambers, Jacobson, Khalil, & Lazzaro, 2019). Despite using two predator species, we did not detect selective predation by either of the hunting spider types (Fig. 2A and B for jumping and wolf spiders, respectively).

**Figure 2.**
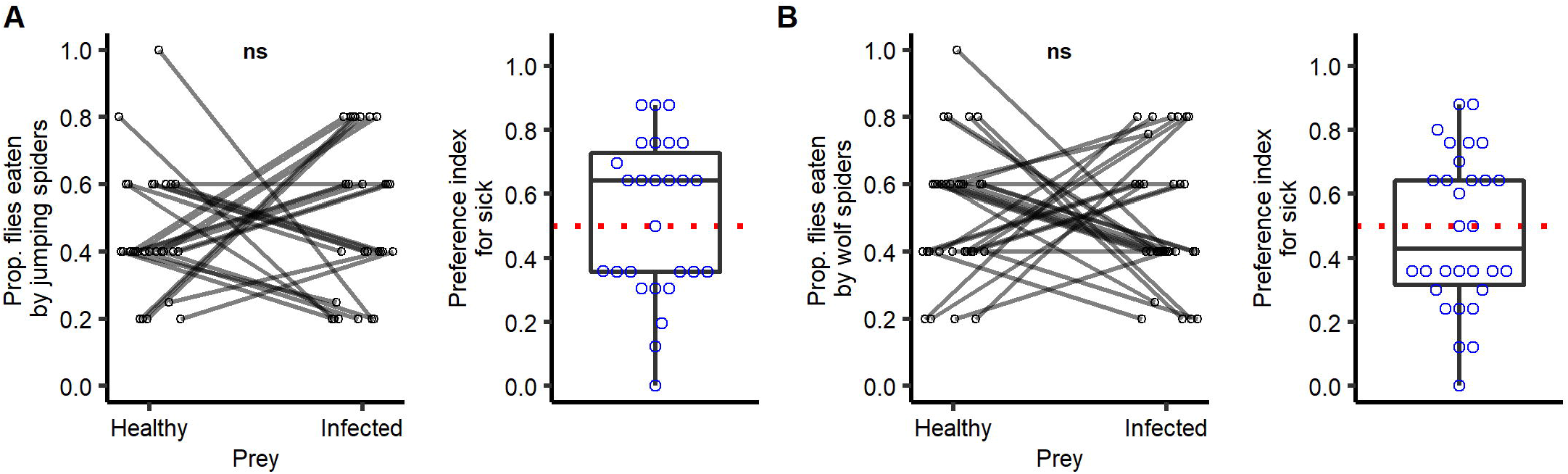
Predation by hunting spiders on a population of *Drosophila melanogaster* Canton S, of either healthy flies or flies chronically infected with *P. rettgeri*. A. Predation by jumping spiders (Hypergeometric test: n= 27 trials, *p*-value > 0.05, Median pref. index = 0.64) B. Predation by wolf spiders (Hypergeometric test: n= 30 trials, *p*-value > 0.05, Median pref. index = 0.43). Each dot represents the proportion of individuals eaten over five flies and the lines connect the groups of individuals from a same trial. Each trial represents ten flies (five from each type) presented to one spider at once. Preference index is calculated as in Manly (1972) and ranges from 0 to 1, from a preference to healthy flies to a preference for sick flies, respectively.

### Predation of flies with cancer by hunting

We asked if cancer of the gut, which we induced, affected the risk for predation of male flies by the hunting spiders. Hyperplasia is the enlargement of an organ or tissue caused by increased cell division and is often an initial stage in the development of cancer. We observed that flies in this early hyperplastic stage (*esg^ts^* > *UAS-Ras^v12^*) did not show an increased likelihood of getting preyed upon by the jumping spiders, which are active diurnal hunters (Fig. 3A). However, jumping spiders did catch more successfully flies with an advanced stage of cancer (*esg^ts^* > *UAS-Ras^v12^*, *UAS-APC-IR*) compared with healthy flies. Furthermore, the preference index for sick individuals increased with the length of time after the tumour had been induced (Fig. 3B).

**Figure 3.**
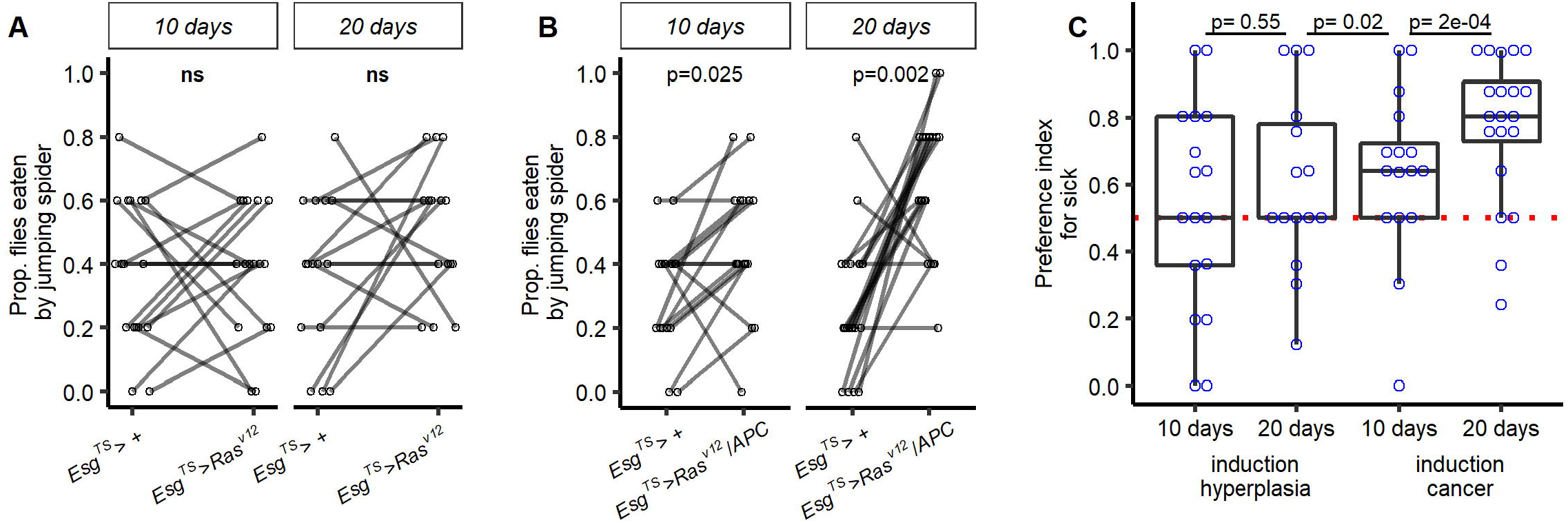
Predation by jumping spiders in a population of *Drosophila melanogaster* that were either healthy or in which colon tumours (hyperplasia) had been induced. Tumour size increased with induction time, during which uncontrolled cell division occurred. A. Uncontrolled cell division was triggered for 10 or 20 days in *esg^TS^* > *Ras^v12^* flies to lead to hyperplasia (respectively with n= 17 and 15 trials). In both cases, individuals with hyperplasia were not differently predated upon compared with the healthy individuals. B. Fast uncontrolled cell division was triggered for 10 or 20 days in *esg^TS^* > *Ras^v12^*, *APC-RNAi* flies to lead to cancer (respectively with n= 16 and 20 trials). In both cases, individuals with cancer were more likely to be predated upon than individuals which did not have cancer. This pattern tended to be stronger as the tumour grew larger (Ordinal logistic regression: LRT=8.07, df=1, p=0.004). Each dot in A and B represents the proportion of individuals eaten over five flies and the lines connect the groups of individuals from a same trial. Each trial represents ten flies (five from each type) presented to one spider at once. Preference index is calculated as in Manly (1972) and ranges from 0 to 1, from a preference to healthy flies to a preference for sick flies, respectively.

## Discussion

Many diseases do not kill their hosts and are considered benign in optimal conditions, such as in a clinic or laboratory. However, it is very likely that non-lethal illnesses still increase the risk of mortality through indirect interactions with other environmental factors. Indeed, being sick may reduce foraging and mating abilities, increase the risk for lethal superinfections, or of being preyed upon. Even though many biological questions are presently being addressed using an integrative approach, the question of whether a non-lethal disease is truly benign in natural conditions seems to have been overlooked in many cases. We hope to generate interest in evaluating the lethality of diseases when their interaction with other risks, such as predation, is considered. Chronic infection by a wild-caught *Drosophila* pathogen did not significantly affect the risk of being preyed upon in the laboratory. In contrast, we observed that while the presence of early stage tumours did not change predation risk, the presence of advanced cancer did compared to age-matched controls.

Little is known about selective predation in individuals with non-infectious diseases. The prevailing opinion that general health, which can be affected by non-infectious sickness, correlates with the risk of being preyed upon, is largely observational (Genovart et al., 2010; Hoey & McCormick, 2004). Nonetheless, although we should not be jumping to hasty conclusions, our results on the effects of non-infectious disease on selective predation suggests a stronger selective pressure than natural chronic infections.

Cancers are common in multicellular organisms and can occur whenever cells evade the normal cell checkpoints that control cell division, proliferation, and apoptosis, thus leading to uncontrolled cell division (Aktipis et al., 2015). Individuals always have some tumours, most of which never become lethal (Abu-Helil & van der Weyden, 2019; Bissell & Hines, 2011). Our understanding of oncogenic phenomena in wild populations of animals is limited, in particular whether tumours that are clinically non-lethal affect the well-being of organisms in the wild (McAloose & Newton, 2009; Pesavento, Agnew, Keel, & Woolard, 2018; M. Vittecoq et al., 2015; Marion Vittecoq et al., 2013). Theoretical modelling further suggests that biotic interactions complicate the predictions regarding the impact of cancer on populations and that prey and predator populations are likely to suffer differently (Perret et al., 2020). If tumour formation is inevitable, it is likely that multicellular organisms have evolved ways to control and tolerate them, perhaps resulting in a trade-off between surviving the tumour and surviving other perils, such as infections or predation (Pavard & Metcalf, 2019). In some cases, selection for tumour tolerance may have arisen from selection for tolerance to other factors, perhaps explaining why early stages of cancer, such as the colon hyperplasia that we induced in *Drosophila*, do not change the likelihood of being preyed upon. In our study, when the tumour developed into a cancer, predation of the flies increased, suggesting that uncontrolled tumours strongly increase the likelihood of being preyed upon. This conclusion supports the idea that individuals with uncontrolled tumours will suffer the risk of increased predation compared with healthy individuals.

Predation is a critical force shaping natural selection (Wade & Kalisz, 1990). By removing prey genotypes susceptible to tumours, predators are likely to have selected for the tumour tolerance largely observed in nature (Bissell & Hines, 2011). Furthermore, by removing individuals with advanced tumours, predation may explain why cancerous animals are rarely recorded in autopsies of wild animal populations (Marion Vittecoq et al., 2013).

We do not know why cancerous flies are more predated than their age-matched controls or even if this result could be to some extend dependent on the sex of the fly. Jumping spiders have been shown to be able to choose prey based on odour and coloration (Vickers & Taylor, 2018). It would be difficult to interpret why such behaviour would have evolved in the context of cancer but we cannot exclude that, spiders recognized and preferred eating cancerous flies. Assays in gut cancerous flies from another study in another genetic background suggests that induction of colon cancer does not affect strongly fly locomotion (Dawson et al., 2018). However, there is still the possibility that cancerous flies have worst reflex than healthy flies or that subtle locomotion differences may be enough to explain a difference in the chance to escape predation. Our results lay the premise of future studies on understanding the mechanisms explaining the reasons for this difference.

The progression of cancer can, in many ways, be compared with infectious diseases. In fact, cancer cells have already been considered as a parasitic species consuming the host’s resources after having emerged from healthy cells (Capp & Thomas, 2020; Duesberg, Mandrioli, McCormack, & Nicholson, 2011). However, if both reduce viability and survival, infectious diseases are, unlike oncogenic processes, under selective pressure to ensure transmission of the pathogen to another host. For both non-infectious diseases and cancer, selective predation could increase the selection against the sickness, as diseases that are moderate and non-lethal could lead to a higher risk of the host being preyed upon, resulting in mortality rates that are similar to those of infectious diseases. With infectious diseases, there may be additional consequences. First, selective predation could affect the transmission of pathogens, spreading them through predator feces over long distances. Second, it might affect epidemics, either preventing them by removing parasite spreaders from the host population (Duffy et al., 2005) or by increasing their likelihood of occurrence by dispersing parasites instead of containing them in dead hosts (Strauss et al., 2016). An additional consideration is that predation may affect the evolution of parasite virulence. It is generally assumed that predation can reduce virulence because hosts with rapidly proliferating parasites would suffer more from the infection and may be eaten before transmitting the parasite to another host, whereas the less rapidly proliferating parasite would impact less host health and would be transmitted before the host is eaten (Moller et al., 2010). However, lower virulence could evolve also through lower pathogenicity, that is by inducing less damage to the host, and not only through reduced proliferation. The most important factor being that bacteria can proliferate within their host as long as the symptoms do not increase the risk of predation. The bacteria studied here, *P. rettgeri*, was obtained from the wild and is prevalent in populations of *D. melanogaster* (Juneja & Lazzaro, 2009). Despite being chronic and the fact that the pathogen can trigger an immune response through the course of the infection (Chambers et al., 2019; Duneau et al., 2017), the chronic infection did not increase the predation by the two species of predator we used in our study. This result supports the hypothesis that chronic infections are selected for such that the symptoms that enhance predation are reduced.

It is reasonable to assume that the success of predators is not simply due to favourable luck. In fact, selective predation may be more the rule than the exception and is likely to have a role on the evolution of diseases, infectious or not (Møller, 2008). On the one hand, selective predation on sick individuals is likely to counter-select against parasites or genetic diseases more strongly than if selection occurred upon disease-mediated host death later in life in absence of predation. On the other hand, in the same way that some parasites evolved to manipulate their host to increase transmission to intermediate hosts (Hughes & Libersat, 2019), parasites may have evolved to make their hosts less conspicuous to predation, for example, by interfering with the search for mates or by lessening the effects of the disease on the host; for example, by reducing symptoms. Because cancer progression generally results in the death of the organism, we cannot expect the incidence of lethal cancer to have evolved in the same way as parasites, except for transmissible cancers such as in Tasmanian devils (Dujon, Bramwell, Roche, Thomas, & Ujvari, 2021). We argue that selective predation on prey with cancer may be one of the reasons why tumours are commonly tolerated in animals, where advanced cancer is only rarely detected.

## Acknowledgements

We thank Violette Chiara, Jean-Baptiste Ferdy, Raphaёl Jeanson, and Daniel Zurek for thoughtful discussion; Pierrick Blanchard, Dieter Ebert, Simon Fellous, Sabine Noebel, and Jennifer Regan for their comments on the manuscript; Cole Gilbert for providing the jumping spiders used in the cancer trials; and Christian Faucher for laboratory support. DD was supported by the French Laboratory of Excellence project ‘TULIP’ (ANR-10-LABX-41; ANR-11-IDEX-0002–02) and the LIA BEEG-B (Laboratoire International Associé-Bioinformatics, Ecology, Evolution, Genomics and Behaviour) (CNRS).

## Author contribution

DD and NB designed and performed the study. DD analysed the data. DD wrote the manuscript with the contribution of NB.

## Data availability statement

All data will be made publicly available.

## References

Abu-Helil, B., & van der Weyden, L. (2019). Metastasis in the wild: investigating metastasis in non-laboratory animals. Clinical and Experimental Metastasis, 36(1), 15–28. doi:10.1007/s10585-019-09956-3

Adelman, J. S., Mayer, C., & Hawley, D. M. (2017). Infection reduces anti-predator behaviors in house finches. Journal of Avian Biology, 48(4), 519–528. doi:10.1111/jav.01058

Aktipis, C. A., Boddy, A. M., Jansen, G., Hibner, U., Hochberg, M. E., Maley, C. C., & Wilkinson, G. S. (2015). Cancer across the tree of life: Cooperation and cheating in multicellularity. Philosophical Transactions of the Royal Society B: Biological Sciences, 370(1673). doi:10.1098/rstb.2014.0219

Albuquerque, T. A. F., Drummond do Val, L., Doherty, A., & de Magalhães, J. P. (2018). From humans to hydra: patterns of cancer across the tree of life. Biological Reviews, 93(3), 1715–1734. doi:10.1111/brv.12415

Arnal, A., Jacqueline, C., Ujvari, B., Leger, L., Moreno, C., Faugere, D.,… Thomas, F. (2017). Cancer brings forward oviposition in the fly Drosophila melanogaster. Ecology and Evolution, 7(1), 272–276. doi:10.1002/ece3.2571

Bissell, M. J., & Hines, W. C. (2011). Why don’t we get more cancer? A proposed role of the microenvironment in restraining cancer progression. Nature Medicine, 17(3), 320–329. doi:10.1038/nm.2328

Brunner, F. S., Anaya-Rojas, J. M., Matthews, B., & Eizaguirre, C. (2017). Experimental evidence that parasites drive eco-evolutionary feedbacks. Proceedings of the National Academy of Sciences, 114(14), 3678–3683. doi:10.1073/pnas.1619147114

Brunner, F. S., Deere, J. A., Egas, M., Eizaguirre, C., & Raeymaekers, J. A. M. (2019). The diversity of eco evolutionary dynamics: Comparing the feedbacks between ecology and evolution across scales. Functional Ecology, 33(1), 7–12. doi:10.1111/1365-2435.13268

Buchon, N., Broderick, N. A., Kuraishi, T., & Lemaitre, B. (2010). *Drosophila* EGFR pathway coordinates stem cell proliferation and gut remodeling following infection. BMC Biology, 8. doi:10.1186/1741-7007-8-152

Capp, J. P., & Thomas, F. (2020). A Similar Speciation Process Relying on Cellular Stochasticity in Microbial and Cancer Cell Populations. IScience, 23(9), 1–12. doi:10.1016/j.isci.2020.101531

Chambers, M. C., Jacobson, E., Khalil, S., & Lazzaro, B. P. (2019). Consequences of chronic bacterial infection in *Drosophila melanogaster*. PloS One, 14(10), e0224440. doi:10.1371/journal.pone.0224440

Chesson, J. (1978). Measuring preference in selective predation. Ecology, 59(2), 211–215. doi:10.2307/1936364

Cunningham, C. X., Johnson, C. N., & Jones, M. E. (2020). A native apex predator limits an invasive mesopredator and protects native prey: Tasmanian devils protecting bandicoots from cats. Ecology Letters, 23(4), 711–721. doi:10.1111/ele.13473

Dawson, E. H., Bailly, T. P. M., Dos Santos, J., Moreno, C., Devilliers, M., Maroni, B.,… Mery, F. (2018). Social environment mediates cancer progression in *Drosophila*. Nature Communications, 9(1), 3574. doi:10.1038/s41467-018-05737-w

DeBlieux, T., & Hoverman, J. (2019). Parasite-induced vulnerability to predation in larval anurans. Diseases of Aquatic Organisms, 135(3), 241–250. doi:10.3354/dao03396

Duesberg, P., Mandrioli, D., McCormack, A., & Nicholson, J. M. (2011). Is carcinogenesis a form of speciation? Cell Cycle, 10(13), 2100–2114. doi:10.4161/cc.10.13.16352

Duffy, M. A., Hall, S. R., Tessier, A. J., & Huebner, M. (2005). Selective predators and their parasitized prey: Are epidemics in zooplankton under top-down control? Limnology and Oceanography, 50(2), 412–420. doi:10.4319/lo.2005.50.2.0412

Dujon, A. M., Aktipis, A., Alix-Panabières, C., Amend, S. R., Boddy, A. M., Brown, J. S.,… Ujvari, B. (2021). Identifying key questions in the ecology and evolution of cancer. Evolutionary Applications, 14(4), 877–892. doi:10.1111/eva.13190

Dujon, A. M., Bramwell, G., Roche, B., Thomas, F., & Ujvari, B. (2021). Transmissible cancers in mammals and bivalves: How many examples are there?: Predictions indicate widespread occurrence. BioEssays, 43(3), 1–10. doi:10.1002/bies.202000222

Duneau, D., Ferdy, J.-B., Revah, J., Kondolf, H. C., Ortiz, G. A., Lazzaro, B. P., & Buchon, N. (2017). Stochastic variation in the initial phase of bacterial infection predicts the probability of survival in *Drosophila melanogaster*. ELife, 6, e28298. doi:10.7554/eLife.28298

Edgar, W. D. (1969). Prey and predators of the Wolf spider Lycosa lugubris. Journal of Zoology, 159(4), 405–411. doi:10.1111/j.1469-7998.1969.tb03897.x

Furey, N. B., Bass, A. L., Miller, K. M., Li, S., Lotto, A. G., Healy, S. J.,… Hinch, S. G. (2021). Infected juvenile salmon can experience increased predation during freshwater migration. Royal Society Open Science, 8(3). doi:10.1098/rsos.201522

Gallagher, S. J., Tornabene, B. J., DeBlieux, T. S., Pochini, K. M., Chislock, M. F., Compton, Z. A.,… Hoverman, J. T. (2019). Healthy but smaller herds: Predators reduce pathogen transmission in an amphibian assemblage. Journal of Animal Ecology, 88(10), 1613–1624. doi:10.1111/1365-2656.13042

Genovart, M., Negre, N., Tavecchia, G., Bistuer, A., Parpal, L., & Oro, D. (2010). The young, the weak and the sick: Evidence of natural selection by predation. PLoS ONE, 5(3), 9774. doi:10.1371/journal.pone.0009774

Gooding, E. L., Kendrick, M. R., Brunson, J. F., Kingsley-Smith, P. R., Fowler, A. E., Frischer, M. E., & Byers, J. E. (2020). Black gill increases the susceptibility of white shrimp, Penaeus setiferus (Linnaeus, 1767), to common estuarine predators. Journal of Experimental Marine Biology and Ecology, 524, 151284. doi:10.1016/j.jembe.2019.151284

Goren, L., & Ben-Ami, F. (2017). To eat or not to eat infected food: a bug’s dilemma. Hydrobiologia, 798(1), 25–32. doi:10.1007/s10750-015-2373-3

Hall, S. R., Cáceres, C. E., Duffy, M. A., & Cáceres, C. E. (2005). Selective predation and productivity jointly drive complex behavior in host-parasite systems. American Naturalist, 165(1), 70–81. doi:10.1086/426601

Hanahan, D., & Weinberg, R. A. (2011). Hallmarks of cancer: The next generation. Cell, 144(5), 646–674. doi:10.1016/j.cell.2011.02.013

Hoey, A. S., & McCormick, M. I. (2004). Selective predation for low body condition at the larval-juvenile transition of a coral reef fish. Oecologia, 139(1), 23–29. doi:10.1007/s00442-004-1489-3

Hollings, T., Jones, M., Mooney, N., & Mccallum, H. (2014). Trophic Cascades Following the Disease-Induced Decline of an Apex Predator, the Tasmanian Devil. Conservation Biology, 28(1), 63–75. doi:10.1111/cobi.12152

Hollings, T., Jones, M., Mooney, N., & McCallum, H. (2016). Disease-induced decline of an apex predator drives invasive dominated states and threatens biodiversity. Ecology, 97(2), 394–405. doi:10.1890/15-0204.1

Holmberg, R. G., & Turnbull, A. L. (1982). Selective predation in a euryphagous invertebrate predator, pardosa vancouveri (Arachnida: Araneae). The Canadian Entomologist, 114(3), 243–257. doi:10.4039/Ent114243-3

Houtz, P., Bonfini, A., Bing, X., & Buchon, N. (2019). Recruitment of adult precursor cells underlies limited repair of the infected larval midgut in *Drosophila*. Cell Host and Microbe, 26(3), 412–425.e5. doi:10.1016/j.chom.2019.08.006

Hughes, D. P., & Libersat, F. (2019). Parasite manipulation of host behavior. Current Biology, 29(2), R45–R47. doi:10.1016/j.cub.2018.12.001

Johnson, P. T. J., Stanton, D. E., Preu, E. R., Forshay, K. J., & Carpenter, S. R. (2006). Dining on disease: How interactions between infection and environment affect predation risk. Ecology, 87(8), 1973–1980. doi:10.1890/0012-9658(2006)87[1973:DODHIB]2.0.CO;2

Juneja, P., & Lazzaro, B. P. (2009). *Providencia sneebia* sp. nov. and *Providencia burhodogranariea* sp. nov., isolated from wild *Drosophila melanogaster*. International Journal of Systematic and Evolutionary Microbiology, 59(5), 1108–1111. doi:10.1099/ijs.0.000117-0

Land, M. F., & Nilsson, D.-E. (2012). Animal eyes. (Oxford University Press, Ed.) (Oxford Ani). Oxford. Retrieved from http://sro.sussex.ac.uk/id/eprint/38226

Lizotte, R. S., & Rovner, J. S. (1988). Nocturnal capture of fireflies by lycosid spiders: visual versus vibratory stimuli. Animal Behaviour, 36(6), 1809–1815. doi:10.1016/S0003-3472(88)80120-9

Manly, B. F. J. (1972). Tables for the analysis of selective predation experiments. Researches on Population Ecology, 14(1), 74–81. doi:10.1007/BF02511186

McAloose, D., & Newton, A. L. (2009). Wildlife cancer: A conservation perspective. Nature Reviews Cancer, 9(7), 517–526. doi:10.1038/nrc2665

Mesa, M. G., Poe, T. P., Gadomski, D. M., & Petersen, J. H. (1994). Are all prey created equal? A review and synthesis of differential predation on prey in substandard condition. Journal of Fish Biology, 45, 81–96. doi:10.1111/j.1095-8649.1994.tb01085.x

Millburn, G. H., Crosby, M. A., Gramates, L. S., Tweedie, S., Gelbart, W., Perrimon, N.,… Baker, P. (2016). Fly Base portals to human disease research using *Drosophila* models. DMM Disease Models and Mechanisms, 9(3), 245–252. doi:10.1242/dmm.023317

Miller, M. W., Swanson, H. M., Wolfe, L. L., Quartarone, F. G., Huwer, S. L., Southwick, C. H., & Lukacs, P. M. (2008). Lions and prions and deer demise. PLoS ONE, 3(12). doi:10.1371/journal.pone.0004019

Mirzoyan, Z., Sollazzo, M., Allocca, M., Valenza, A. M., Grifoni, D., & Bellosta, P. (2019). *Drosophila melanogaster:* A model organism to study cancer. Frontiers in Genetics, 10, 1–16. doi:10.3389/fgene.2019.00051

Møller, A. P. (2008). Interactions between interactions: Predator-prey, parasite-host, and mutualistic interactions. Annals of the New York Academy of Sciences, 1133, 180–186. doi:10.1196/annals.1438.007

Moller, A. P., & Erritzoe, J. (2000). Predation against birds with low immunocompetence. Oecologia, 122(4), 500–504. doi:10.1007/s004420050972

Moller, A. P., Erritzoe, J., & Tottrup, N. J. (2010). Predators and microorganisms of prey: goshawks prefer prey with small uropygial glands. Functional Ecology, 24(3), 608–613. doi:10.1111/j.1365-2435.2009.01671.x

Møller, A. P., & Nielsen, J. T. (2007). Malaria and risk of predation: A comparative study of birds. Ecology, 88(4), 871–881. doi:10.1890/06-0747

Murray, D. L. (2002). Differential body condition and vulnerability to predation in snowshoe hares. Journal of Animal Ecology, 71(4), 614–625. doi:10.1046/j.1365-2656.2002.00632.x

Murray, D. L., Cary, J. R., & Keith, L. B. (2006). Interactive effects of sublethal nematodes and nutritional status on snowshoe hare vulnerability to predation. The Journal of Animal Ecology, 66(2), 250. doi:10.2307/6026

Nagata, T., Koyanagi, M., Tsukamoto, H., Saeki, S., Isono, K., Shichida, Y.,… Terakita, A. (2012). Depth perception from image defocus in a jumping spider. Science, 335(6067), 469–471. doi:10.1126/science.1211667

National Cancer Institute. (2020a). NCI Dictionary of Cancer Terms. Retrieved from https://www.cancer.gov/publications/dictionaries/cancer-terms/def/cancer

National Cancer Institute. (2020b). NCI Dictionary of Cancer Terms. Retrieved from https://www.cancer.gov/publications/dictionaries/cancer-terms/def/hyperplasia

Oppliger, A., Christe, P., & Richner, H. (1996). Clutch size and malaria resistance. Nature, 381(6583), 565–565. doi:10.1038/381565a0

Packer, C., Holt, R. D., Hudson, P. J., Lafferty, K. D., & Dobson, A. P. (2003). Keeping the herds healthy and alert: Implications of predator control for infectious disease. Ecology Letters, 6(9), 797–802. doi:10.1046/j.1461-0248.2003.00500.x

Pavard, S., & Metcalf, C. (2019). Trade-offs between mortality components in life history evolution: the case of cancers. Human Evolutionary Demography, 1–26.

Penteriani, V., Del Mar Delgado, M., Bartolommei, P., Maggio, C., Alonso-Alvarez, C., & Holloway, G. J. (2008). Owls and rabbits: Predation against substandard individuals of an easy prey. Journal of Avian Biology, 39(2), 215–221. doi:10.1111/j.0908-8857.2008.04280.x

Perret, C., Gidoin, C., Ujvari, B., Thomas, F., & Roche, B. (2020). Predation shapes the impact of cancer on population dynamics and the evolution of cancer resistance. Evolutionary Applications, 13(7), 1733–1744. doi:10.1111/eva.12951

Pesavento, P. A., Agnew, D., Keel, M. K., & Woolard, K. D. (2018). Cancer in wildlife: patterns of emergence. Nature Reviews Cancer, 18(10), 646–661. doi:10.1038/s41568-018-0045-0

R Core Team. (2020). R: A language and environment for statistical computing. Vienna, Austria. Retrieved from https://www.r-project.org/

Roche, B., Møller, A. P., DeGregori, J., & Thomas, F. (2017). Cancer in Animals: Reciprocal Feedbacks Between Evolution of Cancer Resistance and Ecosystem Functioning. In Ecology and Evolution of Cancer (pp. 181–191). Elsevier. doi:10.1016/B978-0-12-804310-3.00013-2

Rudrapatna, V. A., Cagan, R. L., & Das, T. K. (2012). *Drosophila* cancer models. Developmental Dynamics, 241(1), 107–118. doi:10.1002/dvdy.22771

Sih, A., Crowley, P., Mcpeek, M., Petranka, J., & Strohmeier, K. (1985). Predation, Competition, and Prey Communities: A Review of Field Experiments. Annual Review of Ecology and Systematics, 16(1), 269–311. doi:10.1146/annurev.es.16.110185.001413

Strauss, A. T., Shocket, M. S., Civitello, D. J., Hite, J. L., Penczykowski, R. M., Duffy, M. A.,… Hall, S. R. (2016). Habitat, predators, and hosts regulate disease in *Daphnia* through direct and indirect pathways. Ecological Monographs, 86(4), 393–411. doi:10.1002/ecm.1222

Temple, S. A. (1987). Do predators always capture substandard individuals disproportionately from prey populations. Ecology, 68(3), 669–674. doi:10.2307/1938472

Vickers, M. E., & Taylor, L. A. (2018). Odor alters color preference in a foraging jumping spider. Behavioral Ecology, 29(4), 833–839. doi:10.1093/beheco/ary068

Villegas, S. N. (2019). One hundred years of *Drosophila* cancer research: No longer in solitude. DMM Disease Models and Mechanisms, 12(4). doi:10.1242/dmm.039032

Vittecoq, M., Ducasse, H., Arnal, A., Møller, A. P., Ujvari, B., Jacqueline, C. B.,… Thomas, F. (2015). Animal behaviour and cancer. Animal Behaviour, 101, 19–26. doi:10.1016/j.anbehav.2014.12.001

Vittecoq, Marion, Roche, B., Daoust, S. P., Ducasse, H., Missé, D., Abadie, J.,… Thomas, F. (2013). Cancer: A missing link in ecosystem functioning? Trends in Ecology and Evolution. doi:10.1016/j.tree.2013.07.005

Wade, M. J., & Kalisz, S. (1990). The causes of natural selection. Evolution, 44(8), 1947–1955. doi:10.1111/j.1558-5646.1990.tb04301.x

Wang, C., Zhao, R., Huang, P., Yang, F., Quan, Z., Xu, N., & Xi, R. (2013). APC loss-induced intestinal tumorigenesis in *Drosophila:* Roles of Ras in Wnt signaling activation and tumor progression. Developmental Biology, 378(2), 122–140. doi:10.1016/j.ydbio.2013.03.020

Woods, G. M., Fox, S., Flies, A. S., Tovar, C. D., Jones, M., Hamede, R.,… Bettiol, S. S. (2018). Two Decades of the Impact of Tasmanian Devil Facial Tumor Disease. Integrative and Comparative Biology, 58(6), 1043–1054. doi:10.1093/icb/icy118

Zurek, D. B., & Nelson, X. J. (2012). Hyperacute motion detection by the lateral eyes of jumping spiders. Vision Research, 66, 26–30. doi:10.1016/j.visres.2012.06.011

Zurek, D. B., Taylor, A. J., Evans, C. S., & Nelson, X. J. (2010). The role of the anterior lateral eyes in the vision-based behaviour of jumping spiders. Journal of Experimental Biology, 213(14), 2372–2378. doi:10.1242/jeb.042382

